# A Combination of Machine Learning and PBPK Modeling Approach for Pharmacokinetics Prediction of Small Molecules in Humans

**DOI:** 10.1101/2023.07.17.549292

**Authors:** Yuelin Li, Zonghu Wang, Yuru Li, Jiewen Du, Xiangrui Gao, Yuanpeng Li, Lipeng Lai

## Abstract

Recently, there has been rapid development in model-induced drug development, which has the potential to reduce animal experiments and accelerate drug discovery. Physiologically based pharmacokinetic (PBPK) and machine learning (ML) models are commonly used in early drug discovery to predict drug properties. However, basic PBPK models require a large number of molecule-specific inputs from in vitro experiments, which hinders the efficiency and accuracy of these models. To address this issue, this paper introduces a new computational platform that combines ML and PBPK models. The platform predicts molecule PK profiles with high accuracy and without the need for experimental data.

This study developed a whole-body PBPK model and ML models of plasma protein unbinding (*f*_*up*_), Caco-2 cell permeability, and total plasma clearance to predict the PK of small molecules. Pharmacokinetic profiles were simulated using a “bottom-up” PBPK modeling approach with ML inputs. Additionally, 40 compounds were used to evaluate the platform’s accuracy. Results showed that the ML-PBPK model predicted the area under the concentration-time curve (AUC) with 62.5% accuracy within a 2-fold range, which was higher than using in vitro inputs with 47.5% accuracy.

The ML-PBPK model platform provides high accuracy in prediction and reduces the number of experiments and time required compared to traditional PBPK approaches. The platform successfully predicts human PK parameters without in vitro and in vivo experiments and can potentially guide early drug discovery and development.

## Introduction

Pharmacokinetics (PK) is a critical aspect of drug development, as it describes the absorption, distribution, metabolism, and excretion(ADME) of compounds in the body. During preclinical stages, lead compounds undergo evaluation for their PK properties through in vitro and in vivo animal experiments. The results of these evaluations can be used to rank compounds or optimize their structures based on the correlation between their physicochemical and PK properties. Moreover, these in vitro and animal PK results can be leveraged to predict human PK phenomena and guide clinical trial design through allometric scaling, compartment models, or PBPK models.

In contrast to traditional PK models with allometric scaling, PBPK models have the ability to predict drug concentrations in plasma and various tissues without the need for animal experiments. As a result, the application of PBPK models has significantly increased in drug discovery and development over the past few years.^1^ Three approaches are commonly used in PBPK model prediction, including “top-down,” “middle-out,” and “bottom-up.”

The top-down approach relies predominantly on observed clinical data, while the middle-out approach combines both in vitro and vivo information to determine unknown or uncertain parameters of the model.^2^ The bottom-up approach, in particular, offers the potential to minimi,:e or replace animal PK studies, as it relies solely on in vitro data for drug-related input parameters. However, IMI-Oral Biopharmaceutics Tools projects show limitations of a “bottom-up” approach in human PK predictions, with only half of the area Under the Concentration-Time Curve (AUC) predictions being within a 2-fold prediction error.^3^ This accuracy may be affected by errors in in-vitro experiments or the accuracy of clearance prediction using in vitro hepatic systems. Such limitations may be overcome by using ML models to predict physicochemical properties directly from structures.

Drug-specific input parameters, such as *f*_*up*_, intrinsic hepatic clearance, and volume of distribution (*V d*_*ss*_), have been well predicted using in silico models. Previously, Doha combined in vitro and ML inputs with a minimal PBPK model and evaluated 240 compounds in rats. ML inputs with fup, LogD, and CL showed only 36.1% of systemic plasma AUC within a 2-fold prediction error.^4^ Vector Group then developed a high-accuracy machine learning-integrated modeling platform, a whole-body PBPK model with an optimized Vdss prediction method.^5^

Several studies have been conducted to optimize the calculation methods of Vdss to improve the accuracy of PBPK predictions. However, it is important to note that clearance also plays a significant role in PK prediction. Clearance, which determines the rate of drug elimination from the body, occurs in the liver, kidney, and bile. Early drug discovery focuses primarily on liver metabolism using in vitro experiments in hepatocytes or microsomes. As a result, several PBPK models only consider hepatic clearance from the in vitro to in vivo exploration (IVIVE) approach. This approach may result in the misprediction of clearance due to the exclusion of certain renal or bile elimination processes. Bowman has reported underprediction of clearance from the IVIVE method, with a 42.2% error rate within a 2-fold margin of error in the microsome system.^6^ This highlights the need for total clearance to improve the accuracy of PBPK modeling in the early discovery stage.

In our study, we have developed a rapid ML-PBPK model platform that enables the simulation of human PK from compound structures. *F*_*up*_, Caco-2 cell permeability, and total plasma clearance for humans were predicted using ML models. These predicted results were then used as input parameters for a whole-body PBPK model encompassing 14 tissues. The prediction accuracy of the platform was evaluated for 40 drugs PK profiles in humans to define its applicability for use in early discovery and clinical phases.

## Materials and Methods

### Data Collection

The human *f*_*up*_ model relies on two data sources: the Watanabe study, which provides data for 2139 compounds, and the Votano study,^7,8^ which provides data for 808 compounds. Overlaps between the two datasets were checked, and compounds were removed if two records had values greater than a 2-fold difference. Compounds with values that differed by less than 2-fold were kept from Watanabe s study due to having more significant figures. Caco-2 cell permeability data for 6083 compounds were collected from public sources.^9–11^The human CLt model used intravenous PK parameters from Lombardo’s study.^12^ Compounds were removed if they had duplicates or invalid SMILES, a molecular weight greater than 900 Da, or where CL was none. Additionally, 40 molecules that overlapped with the experimental data were removed. Finally, we created three datasets: *f*_*fup*_ containing 2062 compounds, Caco-2 containing 6083 compounds, and CLt containing 1215 compounds.

The human plasma PK data for the 40 tests were extracted from previously published papers.^13–50^ All PK data were digitized using the free online tool WebPlotDigitizer.

### ML model building

Compounds SMILES were standardized using the ChEMBL standardizer.^51^ Three different methods (RDKit, Mordred, and PaDEL-Descriptors) were used to calculate descriptors.^52^ These methods generated molecular physicochemical properties for each molecule, resulting in 1826 (Mordred), 1444 (PaDEL), and 208 (RDKit) features used for model construction.

The data sets were split into training, validation, and test sets using random selection at an 8:1:1 ratio. Features with variance values less than 0.05 or those with the same information and a correlation coefficient higher than 0.9 were removed. The Boruta Algorithm^53^ was also used to select significant features in a given data set.

Four common approaches to molecular property prediction were used to build *f*_*up*_, Caco-2, and CLt prediction models. These approaches included Support Vector Machine Regression (SVR), Random Forest (RF), XGBoost (XGB), and Gradient Boost Machine (GBM). In contrast to traditional chemical descriptors, message-passing neural networks (MPNNs) have exhibited advancements in molecular modeling and property prediction. MPNNs are a group of graph convolutional neural networks (GCNs) variants that can learn and aggregate local information of molecules through iterative message-passing iterations. Recently, Yang et al.^54^ have proposed a directed MPNN (D-MPNN) and built the open-source package Chemprop for implementation of D-MPNN. D-MPNN constructs a learned molecular representation by operating on the graph structure of the molecule and passing a message through the edge-dependent neural network. In this study, D-MPNN builds the model based on different datasets and uses RDKit descriptors incorporated into D-MPNN to further improve performance.

The hyperparameters of the regression models were optimized with Bayesian optimization search. Five-fold cross-validation was used to check the stability and predictive ability of the model. Additionally, the performance of the regression models was assessed by the coe cient of determination (*R*^2^) and root-mean-square error (RMSE).

### PBPK model building

Figure 1a shows the compartment model for each tissue, which includes plasma, blood cells, interstitial space, and intracellular space, as previously discussed by Kawai.^55^ Molecules move between adjacent compartments through passive diffusion and connect to the circulatory system through blood ow. The permeability of tissues was assumed to be the same and obtained through Caco-2 cell permeability. Figure 1b illustrates the structure of venous blood, which comprises only plasma and blood cells.

**Figure 1:**
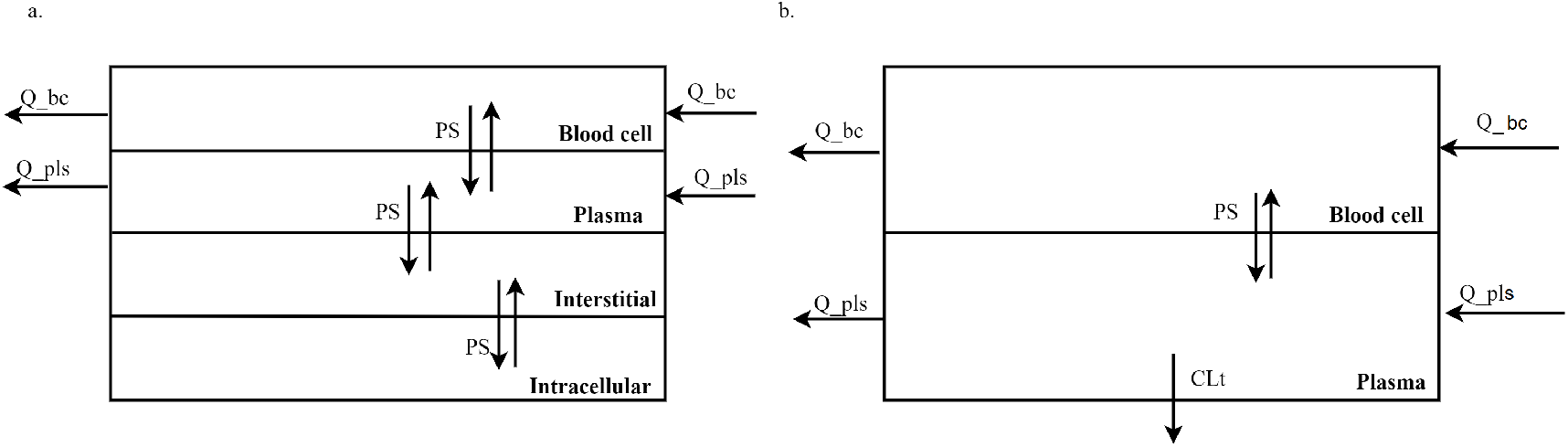
Structure of PBPK model for tissues and venous blood.Each tissue is divided into blood cells, plasma, interstitial and intracellular spaces(a), and each blood vessel only has vascular compartments(b).Details of the model is presented in the method section.

In the ML-PBPK model, three key parameters, fup, caco-2 cell permeability, and CLt, were taken from predictions by ML models, and systemic drug elimination was assumed to occur in venous blood plasma through CLt. On the other hand, in the in vitro input model, a PBPK model using in vitro inputs, all in vitro parameters were taken from experiments. Specifically, elimination processes include hepatic clearance from microsome stability experiments using the IVIVE method and renal clearance as glomerular filtration rate. The differential equations for venous blood vessels are described below as equations 1 and 2. The equations for arterial blood and the portal vein are the same as for venous blood, except for the elimination process.

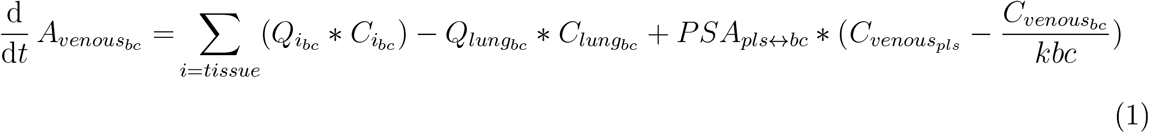

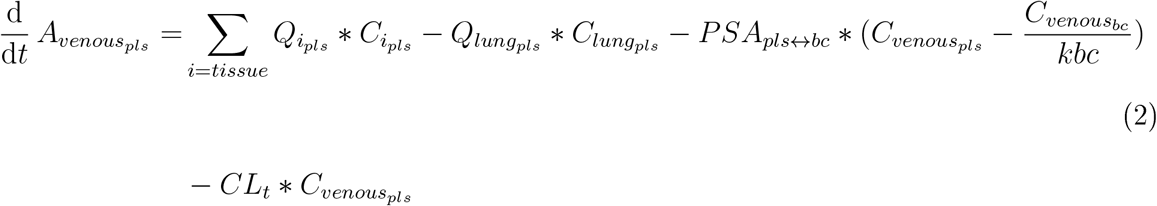

All physiological parameters were adapted from literature,^56–58^ including tissue volumes, blood flow rates, surface areas(SA), tissue compositions, and tissue pH. Tissue partition coefficients (Kp) and blood: plasma ratio (BP) were calculated based on the Rowland-Roger method.^59^ Drug physicochemical properties such as LogP, molecular weight, and pKa values were predicted from structure using ChemAxon. For in vitro testing, drug-related parameters such as Caco-2 cell permeability(P) and microsomal half-life were obtained from previous publications.^60–62^ Physicochemical inputs for the ML-PBPK model simulation were predicted by ML models.

When administered intravenously in a short time, such as a bolus, the maximum concentration (*C*_*max*_) in venous plasma is often reported to over-predict compared to the clinical PK profiles. Prediction errors may be due to the different sampling sites, as clinical samples are usually taken from a peripheral vein in the arm.^63^ To avoid this, the plasma concentration profile in peripheral blood was chosen to evaluate prediction accuracy with observed PK.

### Prediction Performance Assessment

PBPK models were used to simulate concentration-time profiles of tested drugs. Inputs for the models included machine learning and in vitro experimental data. Python was used for model development, and the matplotlib package was used to generate figures.

To evaluate the predicted PK data’s accuracy, non-compartmental analyses were conducted. This involved calculating important parameters such as half-life (*T*_1/2_), area under the curve (*AUC*_0−∞_), clearance (CL), and volume of distribution at steady-state (*V d*_*ss*_) using specific equations. *AUC*_0−∞_ and area under the moment curve (*AUMC*_0−∞_) were calculated using the linear-trapezoidal method. The elimination rate constant (*k*_*el*_) was calculated using the linear regression method. Mean residence time (MRT) was calculated by AUMC/AUC.

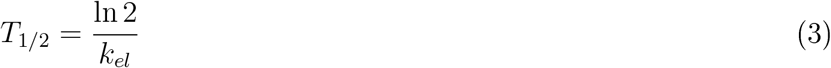

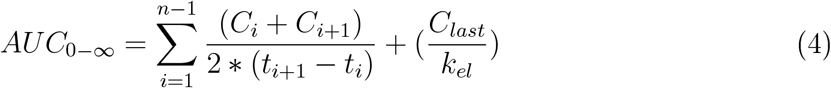

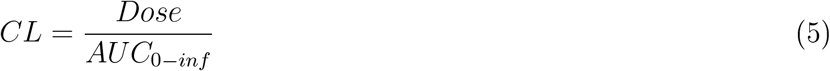

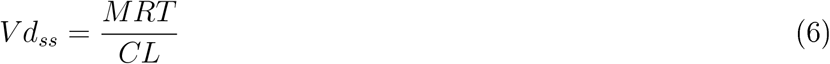

The accuracy of the predicted PK data was measured by calculating the average fold error (AFE) for each PK parameter(Eq. 7). The total number of testing molecules was represented by n. This metric was used to evaluate the overall prediction accuracy of the model. Additionally, the prediction accuracy of the model was assessed by determining the percentage of prediction error within a 2-fold range for each PK parameter.

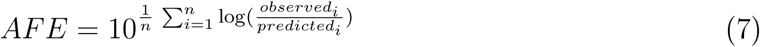

## Results

### ML models

Five ML methods were used to construct models for predicting human *f*_*up*_, Caco-2 cell permeability, and CLt. These methods included SVR, RF, XGB, GBM, and D-MPNN. A series of models were built for each parameter using different training sets. Table 1 presents the statistical evaluation results of the ML models in training and testing.

**Table 1:**
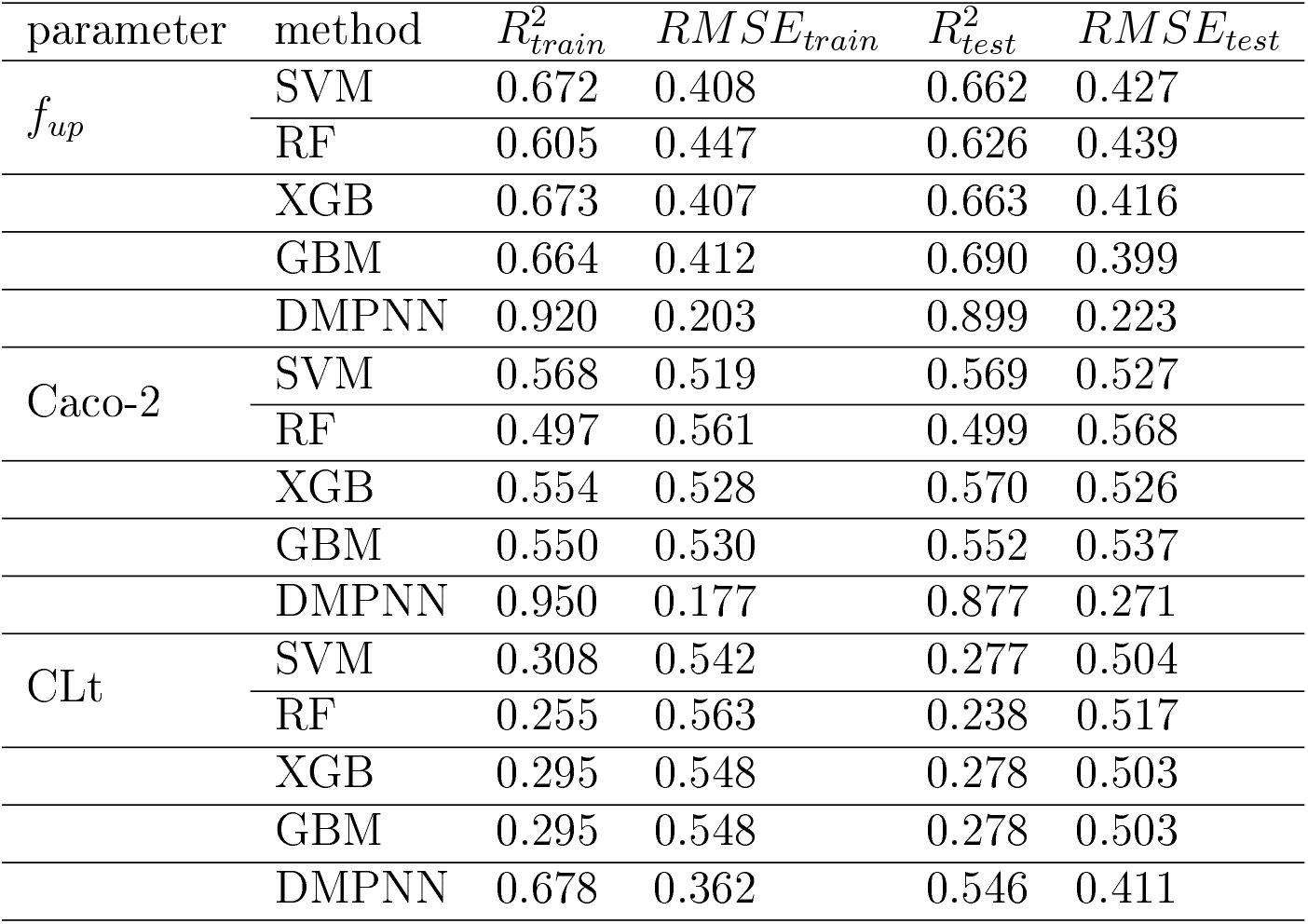
Statistics results of ML model of human *f*_*up*_,Caco-2 cell permeability and CLt.

| parameter | method | $R^2_{train}$ | $RMSE_{train}$ | $R^2_{test}$ | $RMSE_{test}$ |
| --- | --- | --- | --- | --- | --- |
| $f_{up}$ | SVM | 0.672 | 0.408 | 0.662 | 0.427 |
|  | RF | 0.605 | 0.447 | 0.626 | 0.439 |
|  | XGB | 0.673 | 0.407 | 0.663 | 0.416 |
|  | GBM | 0.664 | 0.412 | 0.690 | 0.399 |
|  | DMPNN | 0.920 | 0.203 | 0.899 | 0.223 |
| Caco-2 | SVM | 0.568 | 0.519 | 0.569 | 0.527 |
|  | RF | 0.497 | 0.561 | 0.499 | 0.568 |
|  | XGB | 0.554 | 0.528 | 0.570 | 0.526 |
|  | GBM | 0.550 | 0.530 | 0.552 | 0.537 |
|  | DMPNN | 0.950 | 0.177 | 0.877 | 0.271 |
| CLt | SVM | 0.308 | 0.542 | 0.277 | 0.504 |
|  | RF | 0.255 | 0.563 | 0.238 | 0.517 |
|  | XGB | 0.295 | 0.548 | 0.278 | 0.503 |
|  | GBM | 0.295 | 0.548 | 0.278 | 0.503 |
|  | DMPNN | 0.678 | 0.362 | 0.546 | 0.411 |

The D-MPNN models outperformed the other models for all three parameters(Figure 2). For predicting human *f*_*up*_, the D-MPNN model achieved an *R*^2^ of 0.92 for an independent training set of 2662 compounds and predicted 77.5% (31/40) of the test set within a 2-fold prediction error. The D-MPNN model exhibited the highest *R*^2^ value of 0.95 for the human Caco-2 training set, compared to the GBM model with an *R*^2^ of 0.55. Additionally, the D-MPNN model demonstrated the best predictive ability for human CLt compared to the other models, with 67.5% of the 40 testing compounds predicted within a 2-fold prediction error.

**Figure 2:**
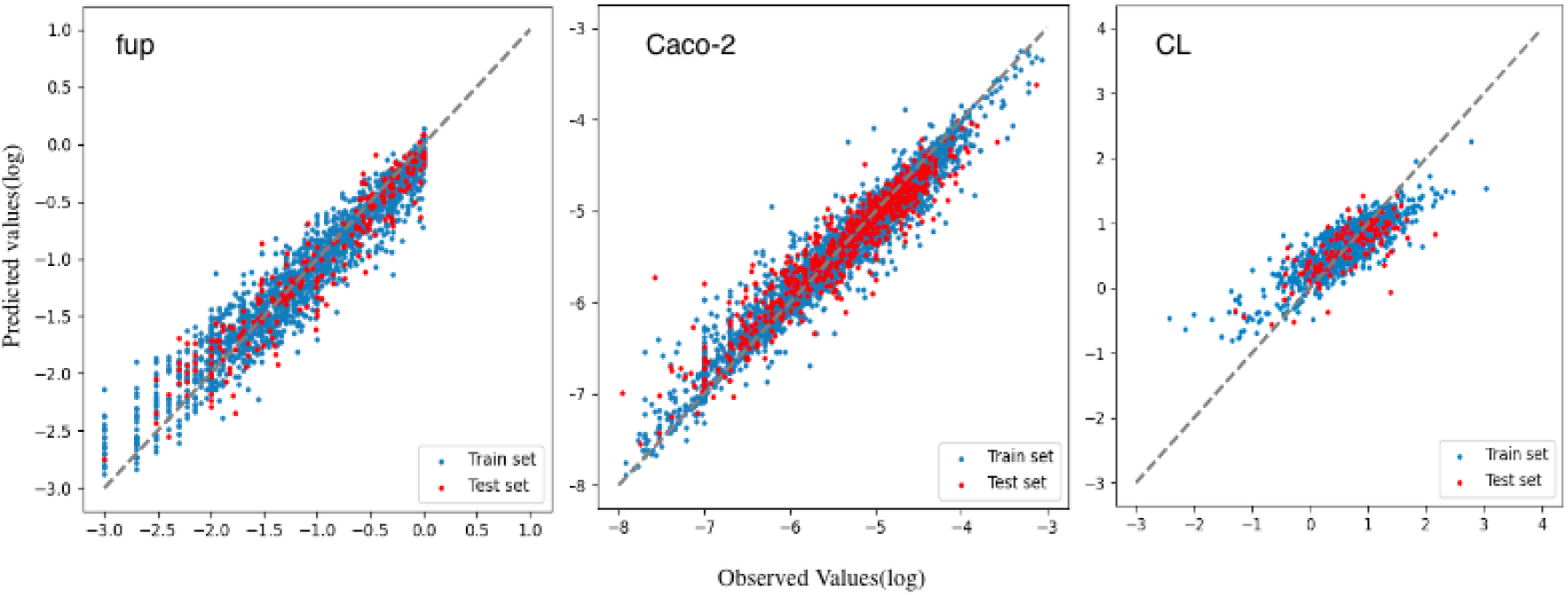
Plots of the observed and predicted *f*_*up*_, Caco-2 cell permeability and CLt of the training set and the test set of the D-MPNN models. The dashed line indicates the line of unity(x=y).

Overall,the best models were used to predict the human *f*_*up*_, Caco-2 cell permeability, and CLt of the forty compounds. These predicted results were then used as inputs in PBPK models.

### PBPK models

Figure 3 compares predicted and observed PK parameters for 40 compounds in humans. All parameters exhibit a good correlation between observed and simulated values, except for CL. Pearson correlation coefficient values (*R*^2^) range from 0.6-0.9. Prediction accuracy of *AUC*_0−∞_ is 62.5% (25/40), with slightly better performance in the ML-PBPK model compared to the in vitro inputs model, which had 47.5% (19/40) within 2-fold error (Table 2). The ML-PBPK model showed relatively good results for CL prediction, with AAFEs of 2.04 and 2.73 and *R*^2^ values of 0.4 and 0.27 respectively, compared to the in vitro inputs model.The predicted/observed ratios of PK parameters show a narrow range with ML inputs (Figure 4), indicating a good agreement between predicted and observed values. Both models showed over- or under-predicted values of CL, with median predicted/observed ratios of 1.37 and 0.77. However, drugs extensively excreted in their unchanged form in urine and with elimination rates higher than normal GFR showed better prediction results with ML inputs, such as Vinorelbine and Atenolol.

**Table 2:** Prediction Accuracy of PK parameters.

| PK parameter | method | AFE | AAFE | %2FE<sup>a</sup> | %3FE<sup>b</sup> |
| --- | --- | --- | --- | --- | --- |
| $T_{1/2}(h)$ | in vitro | 1.44 | 2.22 | 50.0 | 70.0 |
|  | ML | 1.42 | 2.15 | 57.5 | 77.5 |
| $AUC_{0-\infty}(ng * h/ml)$ | in vitro | 1.34 | 2.73 | 47.5 | 62.5 |
|  | ML | 0.800 | 2.04 | 62.5 | 80.0 |
| CL(L/h/kg) | in vitro | 0.750 | 2.73 | 47.5 | 62.5 |
|  | ML | 1.26 | 2.04 | 62.5 | 80.0 |
| $Vd_{ss}(L/kg)$ | in vitro | 0.900 | 2.12 | 57.5 | 72.5 |
|  | ML | 1.31 | 2.12 | 57.5 | 80.0 |

<sup>a</sup> Percentage of prediction error within 2 fold; <sup>b</sup> Percentage of prediction error within 3 fold;

**Figure 3:**
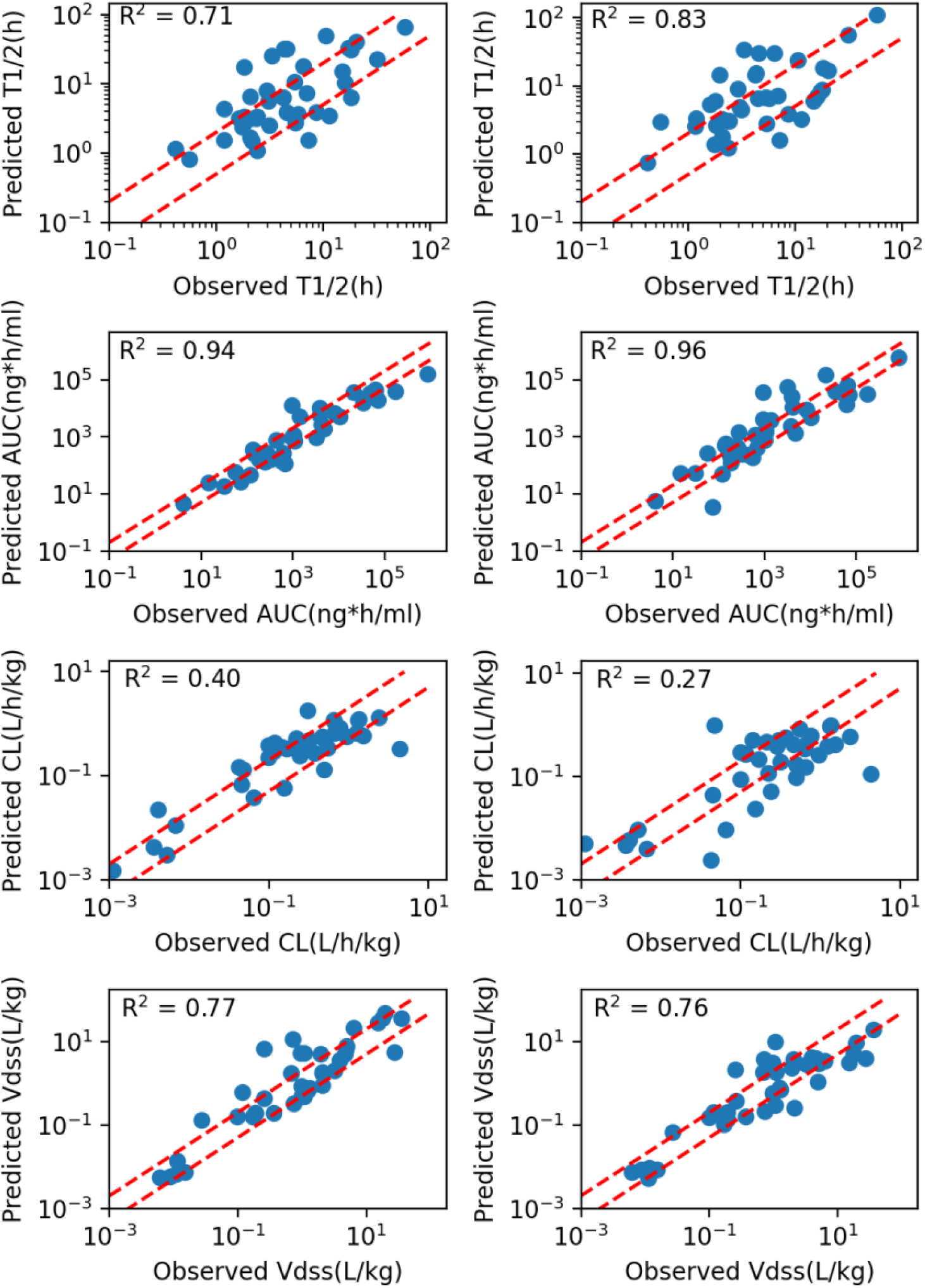
Scatter plots are shown comparision of the predictions and observations for PK parameters after IV dosing in humans using ML inputs (left) and in vitro inputs (right). Two red dashed lines represent±two-fold errors.*R*^2^ were the Pearson correlation coefficient values.

**Figure 4:**
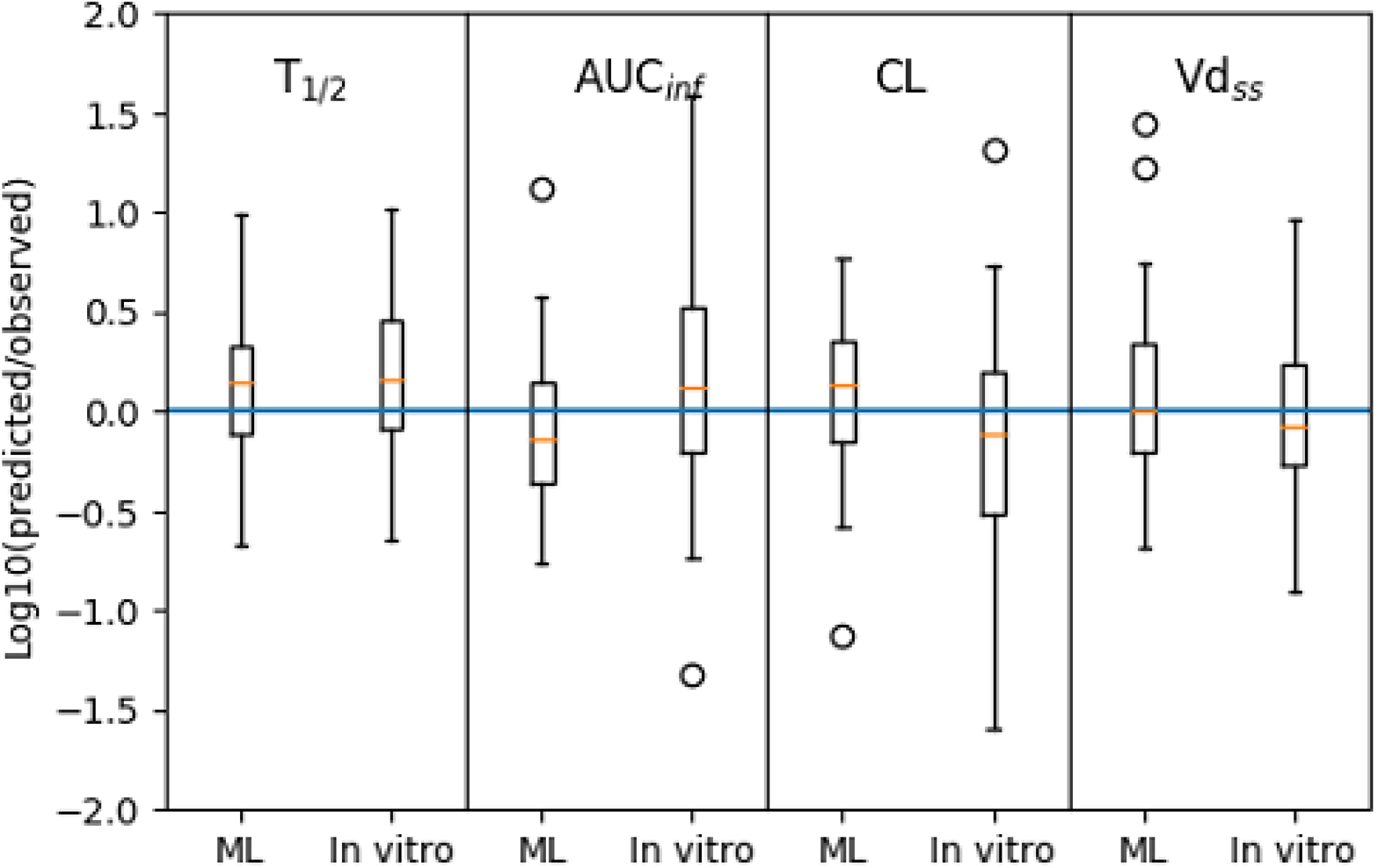
Box plots present log ratios of predicted/observed PK parameter values. Positive values indicate overprediction, and negative values indicate underprediction. The blue line refers to the equal value between prediction and observation.

The results of *V d*_*ss*_ in the ML-PBPK model were similar to those of the in vitro inputs model, as the same tissue partition coefficient calculation method was chosen. *V d*_*ss*_ describes the overall drug distribution in plasma and tissues. In the PBPK model mechanism, drug-related parameters such as *f*_*up*_, cell permeability, and Kp values affect drug distribution into tissues. Since the Kp values for tissues were the same in both the ML-PBPK and in vitro inputs models, comparable *V d*_*ss*_ values indicate that ML prediction of *f*_*up*_ and Caco-2 cell permeability was able to replace experimental values without compromising accuracy. The ML-PBPK model was also more efficient than the in vitro inputs model, with a runtime of only 10 seconds per simulation compared to the few days it may take to collect experimental data and perform model simulations. Overall, the ML-PBPK model was found to be a fast platform that accurately predicts human PK profiles in plasma and tissues.

## Discussion

The development of PBPK models has been a significant advancement in pharmacology. Initially, in vitro data were used to create these models to predict animal and human PK. However, the accuracy of PBPK models and the integrity of input parameters were limited because these measurements did not fully capture the complexity of the human body. As a result, there has been growing interest in developing PBPK models that incorporate inputs without experiments.

One solution is to replace input parameters with ML model predictions. ML has emerged as a powerful tool for predicting drug ADME properties based on their chemical structures. In silico models of physicochemical properties such as *f*_*up*_ and Caco-2 cell permeability have been well studied. Previously, the literature has also focused on improving tissue partition coefficient prediction or hepatic clearance for PK simulation.

In one study by Murad, an ML model was used to predict *V d*_*ss*_ and showed 58% prediction within a 2-fold error.^64^ The Miljkovic group uses ML to predict PK parameters directly from structure, achieving 48.5% within a 2-fold error in their test set.^65^

This study further showed the power of machine learning in predicting relevant parameters that may involve complex physiological processes and hard to be accurately measured by in vitro experiments. A large dataset of drug properties was used to develop ML models that predict *f*_*up*_, Caco-2 cell permeability, and total plasma clearance of drugs. These ML prediction values were then used as inputs for the PBPK models.

Results on 40 drugs showed that the ML-PBPK model predicts human PK parameters with higher accuracy than the in vitro inputs model, and most of the compounds have prediction errors within 2 or 3-fold. Especially, compounds with extensive renal clearance have an average fold error (AFE) of 0.94 compared to the in vitro inputs model of 1.28. In addition, each PK prediction in general was completed within seconds on a machine with the ML-PK model, showing a higher efficiency without compromising accuracy compared to in vitro inputs model.

This study showed how ML methods could improve PK prediction by predicting relevant physicochemical parameters. Future improvements on data quantity and quality that are used to train ML models worth more work. For example, using data from GI organoids may help us train better models to aid the PK prediction for drugs with oral administration. In this study, we predicted total clearance for PK prediction. With more data, we may have separate models for distinct clearance routes, e.g., separate models for hepatic and renal clearance, that provides more information in drug reaserch and development. Thus development of better models as our understanding of deep learning progresses is another valuable direction for future work.

## Conclusions

We evaluated the accuracy of our developed ML-PBPK model platform on 40 compounds by comparing the accuracy of in vitro inputs and ML prediction inputs. The commonly used IVIVE method has limitations in predicting hepatic clearance, and there is a limited experimental exploration of clearance pathways outside the liver in the early stages of drug discovery. As drug clearance is crucial for PK prediction, we used an ML model to predict total human plasma clearance as inputs into the PBPK model for predicting drug concentrations in plasma and tissues. This method was able to guide the development and prioritization of lead compounds based on molecular structure for PK prediction before in vitro experiments. In the future, the accuracy of the ML-PBPK model can be further improved or optimized for specific molecular structures by expanding the training set. Methods such as graph-based multi-task learning, pre-trained models, and model ensembles will be employed to improve accuracy. Furthermore, studying the interpretability of the prediction results is also essential.

## TOC Graphic

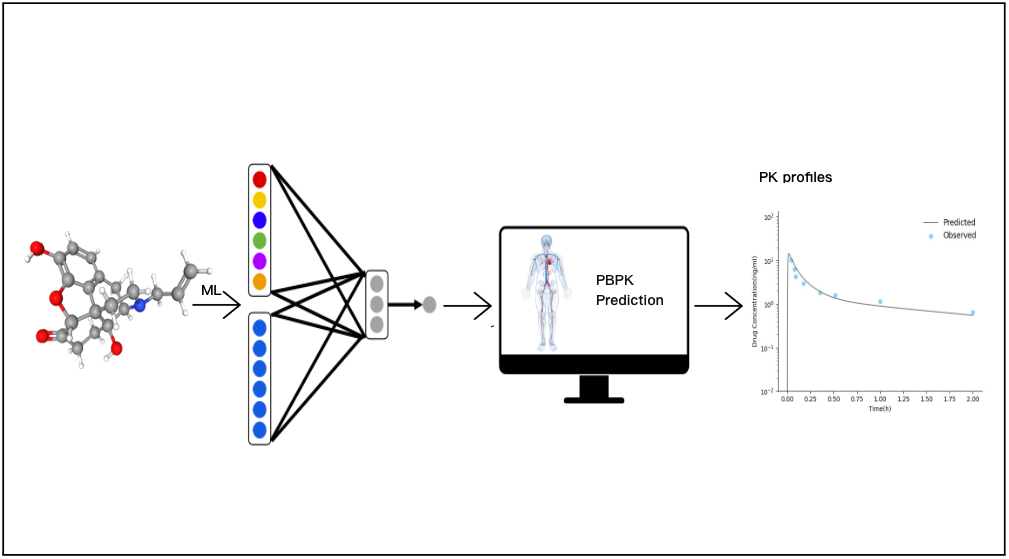

